# Sampling Techniques and Genomic Analysis of Biological Material from Artworks

**DOI:** 10.1101/2024.04.23.589986

**Authors:** Rhonda K. Roby, Rosana A. Wiscovitch-Russo, Rebecca Hart, Amanda E. Appel, Manija A. Kazmi, Thomas Huber, Karina C. Åberg, Thomas P. Sakmar, José A. Lorente, Norberto Gonzalez-Juarbe

## Abstract

The genomic analysis of biological material obtained from artwork or cultural heritage artifacts can be used to guide curation, preservation and restoration. However, the recovery of biological samples from artworks is dependent on the sampling technique used and the media from which the biological materials are recovered. The ideal sampling method should be non-invasive, yet robust. Only rare examples of direct comparisons among sampling techniques have been reported. To develop a framework for the recovery of biological samples for subsequent microbial DNA analysis, we studied five artworks on paper and compared three sampling methods, each with increasing degrees of invasiveness. Minimally invasive swabbing techniques collect samples from the surface, whereas more aggressive techniques such as wet vacuuming were expected to yield more biological material from within the support media. We report a comparison of collection techniques to generate microbial DNA sequence data from artworks on paper. We observed that wet vacuuming resulted in higher DNA recovery than double swabbing and core punches; however, all three techniques yielded satisfactory microbial DNA sequences. Diverse microbial populations existed to varying degrees on the corners and centers of the five artworks studied, but the distribution of the total biomass was relatively even across the surfaces of the works sampled. Studies of peripheral regions, where sampling is less likely to cause alterations to the artwork, could thus yield useful results in microbiome studies. These results provide a framework for sampling artworks on paper to obtain biological material for microbial genomic analysis. The methods described may provide microbiome identification to facilitate restoration and preservation, and might also contribute to the determination of provenance.

## 1. INTRODUCTION

Forensic analysis of artworks is used extensively to gain insights into methodology, technique and provenance [1], and to facilitate restoration efforts [2]. However, most scientific analyses of artwork have focused on direct physical examination using spectroscopic or analytical chemical techniques [2–6]. Advances in molecular biology have facilitated the development of approaches to study ancient microbial and human DNA sampled from archeological sites and burial sites [7–9] as well as environmental DNA (eDNA) [10]. These studies have provided significant insight regarding the genetics of human speciation, global migration, and health history. In principle, similar tools could also be used for the analysis of microbial (or human) DNA sampled from artworks to facilitate restoration and preservation, as well as provenance and attribution. However, systematic studies of DNA sampling and analysis from artworks have not been extensively reported. Some examples of sampling technique publications include a microbial biofilm [11] and a comparison study with swabs, scraping, eraser rubbings, and micro-aspiration [12].

Our overall aim was to advance methodologies to facilitate sampling of DNA from artworks or other cultural/historical artifacts. We compared three sampling techniques to recover biological material suitable for microbial DNA sequencing from five authentic metalpoint and chalk drawings on laid paper. We first compared the yields of DNA from samples obtained using three sampling methods with increasing degrees of invasiveness: swabbing with dry or wet cotton applicators, wet vacuuming with a commercial device, and core punch biopsies. The highest yield of DNA was obtained from the wet vacuum technique. However, the double swab technique and the core punches also resulted in satisfactory microbial DNA yields. The 16S rRNA gene, specific for targeting and amplifying bacterial DNA from the isolated genomic DNA (gDNA) of low biomass samples, was sequenced. In general, the abundance of microbial DNA did not vary significantly among samples taken from the corners, edges, or centers of the artworks. However, differences were observed in the microbiomes among different drawings, as well as in different locations within the same drawing. For example, 150 different operational taxonomic units (OTUs) were found, but only 12 OTUs were shared among all samples collected. Our results provide a methodological framework for collecting samples from artworks on paper to obtain biological material for microbial genomic DNA analysis.

## 2. MATERIALS AND METHODS

### 2.1 Artwork

Since our aim is to develop and benchmark minimally-invasive and non-destructive sampling techniques and compare them with more invasive and potentially-destructive methods, we focused on centuries-old artworks on paper that were judged to hold limited historical or cultural significance as they might be damaged or destroyed in the course of the work. The five artworks used in this study were acquired from reputable e-commerce platforms or from private collections and to our knowledge had not been recently displayed. All works were obtained unframed and were stored in a portfolio folder with plastic sleeve separators at room temperature in the dark. The artworks studied included two metalpoint and three chalk drawings. Metadata (if known) is provided for each piece including the artist’s name and time period, the support matrix and medium, and the size and paper thickness (Table 1). Prior to sampling, high-resolution digital photographs were recorded of the front and back of each drawing using a Sony ILCA- 77M2 camera in raw format (6000 × 4000 pixels) with a Sigma 35-mm f/1.4 DG HSM Art Lens (SIGMA, Kanagawa, Japan) under a variety of lighting conditions using a Bower SFD-RL71 Large Digital LED Ring Light (S. Bower, Inc., Queens, NY, USA). Images were scaled in Adobe Photoshop 2022 (Adobe, San Jose, CA, USA) based on a ruler photographed together with the artwork. A lightbox (LitEnergy A4S LED Copy Board Light Tracing Box, LitEnergy, Guangdong, China) was particularly useful for backlighting and highlighting surface irregularities, textures, watermarks, and translucent elements within the artwork.

**TABLE 1.**
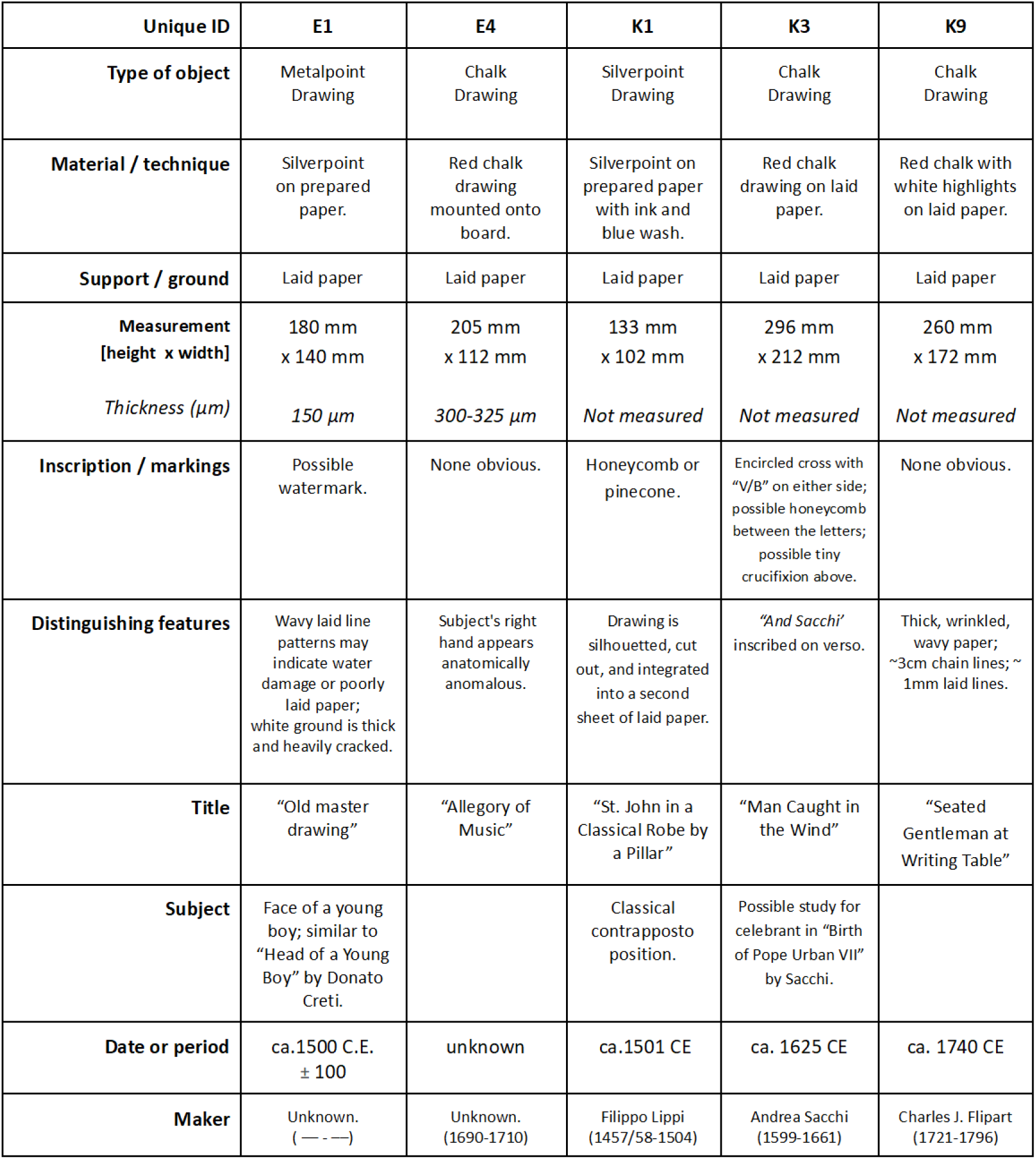
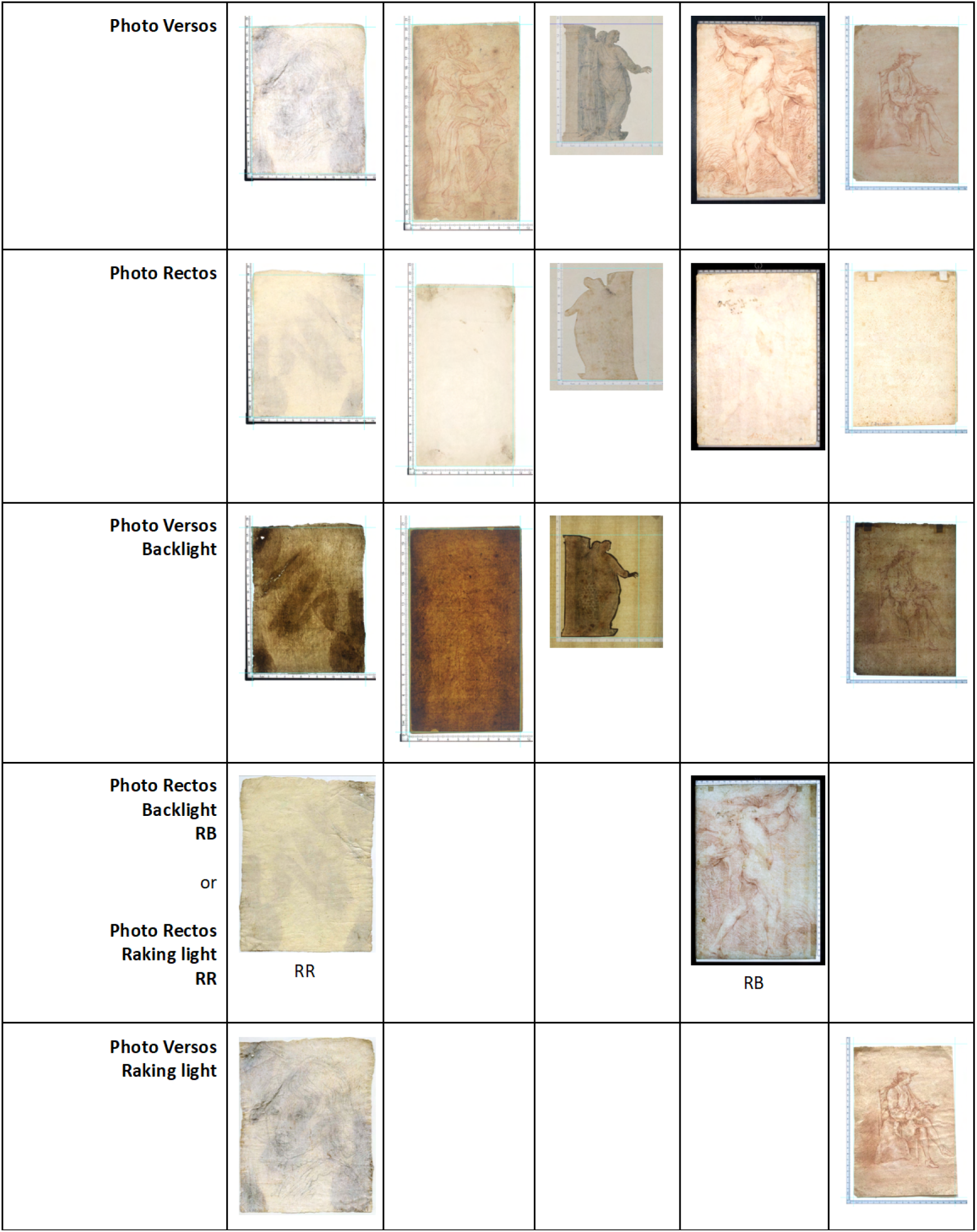
Artworks used in study.

### 2.2 General procedures

Biological samples were collected using sterile laboratory supplies. The use of sterile techniques and personal protective equipment with frequent glove changes minimized the risk of introducing external biological material and/or other external contaminants to the artworks. Bench surfaces and any reusable laboratory supplies or tools were cleaned before each individual work was sampled. A three-step cleaning procedure was used: a 10% diluted commercial sodium hypochlorite solution followed by sterile water rinses and finally ethanol to facilitate the drying of the materials. A fresh fluid barrier paper sheet (Polyshield® Fluid Barrier, VWR, Radnor, PA, or BenchGuard™, Thomas Scientific, Swedesboro, NJ) was used with the impermeable polyethylene coating side up for each sample collection. Sampling controls were used to account for any external factors outside of the artworks that might influence the results.

### 2.3 Alteration scale

When sampling the artworks, a subjective scale of potential alteration to the artwork was employed. An alteration scale provides a systematic approach to categorize, classify, or describe changes, modifications, or variations in the artworks caused by the sampling procedure. Each sampling technique has its own limitations for recovery of biological material along with the potential to damage or alter the artwork being sampled. The alteration scale introduced in this study is designed to quantify and categorize the extent of alternation to an area of the artwork due to the sampling technique used. An alteration score (AS) is assigned for each instance of sampling, and while subjective, provides a framework to document the potential effects of sampling protocols. While art conservators have developed numerous approaches to document damage or decay to artwork [13], this scale is designed to be used specifically to record the effects of techniques used for sampling for biological analysis. The AS scale ranges from AS0 (observation only; no sampling) to AS9 (extensive alteration).

**AS0 (no alteration).** The AS0 rating is assigned when artwork is examined, assessed, and photographed, but where no sampling has taken place. **AS1–AS3 (minimal alteration).** AS1 is assigned when the sampling method used was merely light, dry swabbing of an artwork with solid substrate (*e.g.,* stone, plaster, wood). AS1 is also assigned when the artist’s work area is not directly sampled, for example when swabbing the edge or back of canvas, linen, or paper supports. In some special cases, an area of artwork with certain media such as oil, may be lightly swabbed with no risk of any alteration to the artwork. AS2 to AS3 is assigned when an area of the artist’s work is sampled, but where there is minimal potential alteration. This may involve samples taken by rubbing with a sterile swab, which causes no or minimal potential transfer of the art medium onto the swab. An example of an AS3 rating may be a rubbing of a chalk drawing, which results in the potential for chalk transfer to the swab, but with no alteration of the artwork visible to the unaided eye. **AS4–AS6 (moderate alteration).** A rating in this range indicates that liquids were used during the sampling process, and/or there is visible transfer of the art medium onto the collection device. Even when alteration is inadvertent, it should still be documented. An AS4 to AS6 rating suggests that the artwork touched by the sampling materials was visibly altered. **AS7–AS9 (extensive alteration).** Alterations to artworks in this category have been either wetted by the collection technique or were sampled by core punches, or destructive removal of a portion of the artwork, leading to obvious damage to both the support and the art medium. AS7 to AS9 results in more significant and potentially irreparable changes to the artwork.

The AS serves as an essential record to assess the potential impact of the sampling process on the artwork’s integrity. It can be used to record the direct effects of sampling and should help art conservators, researchers, and historians understand the level of risk associated with various sampling techniques and their effects on the artwork’s condition. By documenting the AS and the potential sampling technique to be used, the trade-off between the need for analysis and the preservation of the artwork’s original state can be evaluated.

### 2.5 Sampling

A systematic approach to sampling similar areas of the artworks using different sampling techniques was defined. Areas to be sampled where defined using virtual triangular zones on the artwork for the purpose of collecting samples from predefined regions. Each of the triangular zones was assigned a letter label. When orienting the artwork for display, the two upper corners (left and right) and an equal area of the artwork in the center were sampled. Using a template of precut plastic and the natural right angle of the artwork, an isosceles right triangle was drawn onto the artwork to guide the prototype sampling. For actual sampling procedures, a template can be drawn onto an acetate film sheet placed between a lightbox and the transilluminated drawing. Two equal isosceles right triangles (together making a square) were used to define the sample area of the center of the artwork. The upper left corner’s isosceles right triangle was divided in half to create two smaller right triangles. The leftmost of the two triangular regions from the upper left was labeled “A” and the rightmost triangular portion was labeled “B.” The upper right’s corner isosceles right triangle was labeled “C.” The square in the center of the artwork was divided into two right isosceles triangles, divided from the upper left corner to the lower right corner. The lower isosceles right triangle was divided in half to create two smaller right triangles “D” and “E.” The upper isosceles right triangle is labeled “F.” (Figure 1). Triangles A, B, D, and E were measured to be 4.25 cm × 4.25 cm × 6 cm (perimeter = 14.5 cm; area = 9 cm^2^); triangles C and F were measured to be 6 cm × 6 cm × 8.5 cm (perimeter = 20.5 cm; area = 18 cm^2^). This geometric approach to sampling facilitates systematic collection of samples from various regions of the artwork, including the upper corners, the central area, and smaller regions within the central square for the different sampling techniques. The use of labels for each triangle ensures that the collected samples can be properly identified and associated with specific areas on the artwork for documentation and further analysis. Due to the different sizes of the artworks, if this configuration was not possible, then a reasonable version of it was applied.

**FIGURE 1.**
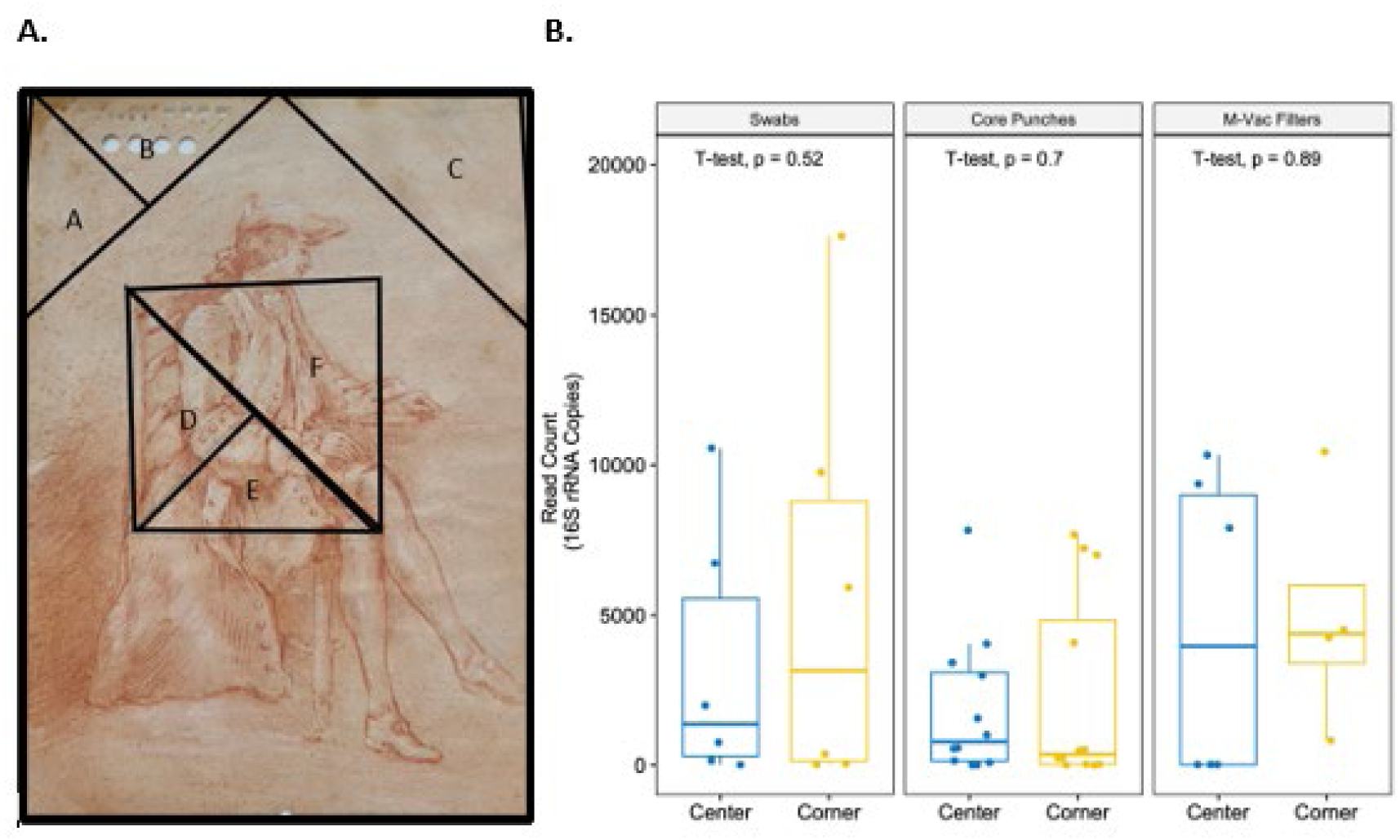
K9 artwork and the geometric areas of sampling and bacterial 16S rRNA read counts. A) Example of an artwork and the designated locations (triangles A, B, C, D, E, and F) for sampling techniques. Swabs were collected from triangles A (corner) and D (center). Core punches were collected from triangles B (corner) and E (center). M-Vac filters were collected from triangles C (corner) and F (center). B) Total read counts (full-length 16S rRNA) of the samples collected using different sampling techniques from the locations noted in panel **A**. There was no significant difference of the mean bacterial DNA read counts of the samples collected from the center or corners of the artwork (t-test).

The collection techniques employed included surface swabbing, core punches taken with a disposable biopsy punch that cuts through the solid support and media, and wet vacuuming the surface, which theoretically can extract samples from within the solid support and the media. Swabbing involves the use of sterile swabs to gently rub and collect samples from the surfaces of artworks. This relatively non-invasive technique allows for the retrieval of surface materials, potentially including biological samples. The use of disposable biopsy punches allows for the core punches of the artwork, both support and media. This technique is suitable for extracting samples from deeper layers or within the solid support material itself. Wet vacuuming is a specialized technique employed to extract samples from the solid support and media of the artworks. It theoretically has the capacity to collect material not only from the surface, but also from within the solid support and media. While wet vacuuming retrieves materials from deeper layers of the support, the stratigraphy of those biological materials will be lost. The nomenclature used for each sampling is described in Table 2.

**TABLE 2.** Nomenclature of sampling from each triangular area.

| Nomenclature |  |  |  |  |  |
| --- | --- | --- | --- | --- | --- |
| Triangles A and D | Swabs | Triangles B and E | Core punches | Triangles C and F | M-Vac filters |
| 1 | Front of artwork | 1 | 1.2 mm Harris punch | a, b, or c | Filter quadrants |
| 2 | Back of artwork | 2 | 2 mm Integra® Miltex® punch | a1 | Filter quadrant, flash freeze |
| a | Wet swab | 3 | 4 mm Integra® Miltex® punch |  |  |
| b | Dry swab | 4 | 6 mm Integra® Miltex® punch |  |  |
| Pu-RB | Puritan swab reagent blank | a | Punch 1 |  |  |
|  |  | b | Punch 2 |  |  |
|  |  | c | Punch 3 |  |  |

#### 2.5.1 Swabbing Techniques

Two techniques have been used most commonly to collect samples from artworks: the “direct swab technique” [14, 15] and the “double-swabbing technique” [16,17]. For the “direct swab technique,” a sterile swab is gently rubbed across the surface of the area to be sampled while swirling it for coverage around the entire swab. One swab is retained as a negative control. The swabs are placed in individual sterile tubes for storage. For the “double-swabbing technique,” a drop of molecular biology grade water, or approximately 50 µL, is added to one swab. The surface of the area to be sampled is rubbed with the swab while swirling. Then, the same surface area is sampled with a second dry swab using the same technique. After air drying, under conditions to avoid environmental contamination, both sampling swabs are placed in a sterile tube for storage. As a control, a drop of water is added to a swab and a second dry swab is also included. Depending on the artworks tested and the regions sampled, direct swab techniques are considered to be AS1 to AS3, and the double-swabbing technique, which involves the application of water to the artwork, is generally considered to be AS4 to AS6, indicating a higher degree of potential alteration.

#### 2.5.2 Core Punches

The collection of core punches from artworks is a destructive technique that removes samples from specific areas for analysis, including both the surface of the artwork (*i.e.*, the art medium) and from the support (*i.e.,* paper, canvas, wood support). Core punches in this study were hypothesized to contain all available biological material that could be potentially extracted from a given full-thickness area and were used as controls. Supplies needed to obtain core-punch samples include the 1.2-mm Harris Uni-Core punch (MilliporeSigma, Burlington, MA) or the disposable biopsy punches (2-mm, 4-mm, or 6-mm diameters) (Integra LifeSciences Production Corporation, Mansfield, MA). The targeted area is identified, the artwork is placed on the barrier, and even pressure is applied at a right angle. While stabilizing the artwork with the free hand, pressure is used to push downward into the artwork while rocking the punch with a semi- circular rotation until the tip passes all the way through the artwork’s support. After coring is complete, the punch/cutter tip is placed over a sterile storage tube and the plunger is depressed to drop the sample into the tube. For some samples, sterile PEEK tubing (readily available in the laboratory) (Amersham Biosciences AB, Uppsala, Sweden) was used to push the cored sample into the sterile storage tube. For quality control, collection of samples on the edges or folds of the artwork were collected to serve as a background control. Core punches were classified as AS7 to AS9, depending on the punched area of the artwork (whether central or peripheral) and the size of the punch. For example, a punch on the periphery of the artwork may be assigned AS7, but the same size punch in the center of the art should be assigned AS9, the maximum AS possible.

#### 2.5.3 Wet vacuum sampling

The M-Vac^®^ System (M-Vac Systems, Inc., Sandy, UT) [18] is a wet-vacuum system that dispenses sterile Surface Rinse Solution (SRS) (M-Vac Systems, Inc.) onto the artwork and uses a vacuum mechanism from a sampling head to collect not only the SRS, but also any suspended cellular material. This technique is advantageous for sampling of larger areas and from various substrates. With the fluid barrier’s impermeable polyethylene coating side up, its surface was cleaned with 70% ethanol and then allowed to air dry. The artwork was placed on the fluid barrier and the M-Vac instrument was powered on. The vacuum operator collects biological samples using a unidirectional sampling method. After the collection of the SRS containing potential biological material, a Nalgene 250-mL filter unit fitted with a 0.45-µm PES filter disc was connected to a vacuum pump. The SRS is slowly poured into the middle of the filter unit with the vacuum pump powered on. The filtered SRS is then poured back into the sample collection bottle, swirled to suspend any cellular material on the sides of the bottle, and filtered again. The use of double filtration enhances the concentration and purity of the collected cellular material. The Nalgene 0.45-µm PES filter is allowed to dry in a laminar flow hood with the lid to the Nalgene unit slightly ajar, ensuring a controlled and sterile environment. Once the filter is fully dry, the filter disc is carefully removed using a sterile scalpel and clean tweezers. The filter is then cut into quadrants, 3 quadrants are placed in a single sterile tube and stored at room temperature and the fourth quadrant is placed in a separate sterile tube and subjected to flash freeze (Speer, 2022) and stored at -80°C. Depending on the artworks tested and the region sampled, an AS7 to AS9 was assigned for the M-Vac wet vacuuming technique. There is undoubtedly alteration to the artwork merely by adding liquid directly onto the solid support and media.

### 2.6 Nomenclature

The sampling nomenclature is described for each sample location (Table 2). Triangles A and D were subjected to collection of biological materials by the double-swabbing technique. For example, sample A1a represents the swab from Triangle A (upper left corner), on the front of the artwork, and the wet swab. Triangles B and E were subjected to core punches using different sized biopsy punches. Sample E3b represents the core punch from Triangle E (center of the artwork), using the 4-mm Integra Miltex punch, and the second of three core punches collected. Triangles C and F were subjected to the collection of biological materials by the wet vacuum system. Cc is the sample obtained from the wet vacuuming from Triangle C (upper right corner on the front of the artwork) and the quadrant c of the collected air-dried filter.

### 2.7 DNA isolation and library preparation

#### 2.7.1 Cleaning and preparing workstation

To avoid contamination of ancient samples with modern human or environmental bacteria DNA during sample processing, DNA isolation and library preparation were conducted in a disinfected vertical laminar flow hood solely designated for processing ancient artwork. The hood and its contents (including pipettes, tip boxes, and racks) were disinfected with 70% ethanol, treated with ELIMinase (Catalog number 1101, Decon Laboratories, Inc., King of Prussia, PA, USA), and then exposed to UV light for 30 minutes. Disinfection and cleaning of the hood was conducted routinely before and after sample processing to maintain sterility and minimize contamination. Additionally, personal protective equipment (*e.g*., laboratory coats, gloves, face masks) were used along with disposable sleeves. Disposable sleeves and gloves were routinely changed during the course of the procedure.

#### 2.7.2 DNA isolation of sample collected from aging artwork

The gDNA was extracted from the swabs, core punches, and M-Vac filters using DNeasy^®^ PowerSoil® Pro Kit (Catalog number 47016, QIAGEN, Hilden, Germany) with some minor modification to steps 1-4 of the manufacturer’s protocol (DNeasy PowerSoil Pro Kit Handbook 06/2023). CD1 buffer (800 μL) was added to the sample tube and incubated at 4°C overnight to ensure lysis of any cells present and preservation of the DNA. The following morning, 800 μL of the sample lysate was collected and transferred into the provided PowerBead Pro Tube and mechanically lysed using PowerLyzer^®^ 24 Homogenizer (110/220 V) (Catalog number 13155, QIAGEN, Hilden, Germany) at 3000 rpm for 30 seconds. Then, DNA extractions were performed according to the manufacturer’s protocol from steps 5-17. As suggested by the protocol, DNA was eluted in a minimum volume (50 μL) of pre-heated C6 buffer (10 mM Tris-HCl) at 65°C. Samples were quantified using Qubit™ 1X dsDNA High Sensitivity Assay Kit (Catalog number Q33231, Thermo Fisher Scientific, Waltham, MA, USA). DNA quantity and quality were also measured using the Nanodrop ND-1000 Spectrophotometer (Thermo Fisher).

#### 2.7.3 Full length 16S (V1-V9) library preparation and sequencing

Libraries were prepared using a tailed primer approach with 16S rRNA gene-specific primers spanning the V1-V9 variable regions. We used forward primer 27F 5’- AGRGTTYGATYMTGGCTCAG-3’ and reverse primer 1492R 5’- RGYTACCTTGTTACGACTT-3’, each configured with 16-bp index sequences and pad regions as per Pacific Biosciences specifications for internal indexing of sample-specific libraries [19]. Using Q5^®^ High-Fidelity 2X Master Mix (Catalog number M0492, New England BioLab, Ipswich, MA, USA), libraries were generated at 50 μL with thermal cycler conditions: 95°C for 3 minutes initial denature, 35 cycles of 95°C for 30 seconds denature, 54°C for 30 seconds annealing, and 72°C for 1 minute extension. PCR products underwent fragment size verification using LabChip® GX Touch™ Nucleic Acid Analyzer (PerkinElmer) and 1% agarose gel electrophoresis. To remove primer dimers (PCR products <100 bp), samples were subjected to bead cleanup using 0.6X SPRIselect (Catalog number B23319, Beckman Coulter, Brea, CA, USA). Libraries were pooled in an equimolar approach prior to construction of SMRTbell® Prep Kit 3.0 (Catalog number 102-182-700, PacBio, Menlo Park, CA, USA) sequencing library preparation according to the manufacturer’s protocol. Sequencing was performed on a PacBio Sequel II/IIe system using SMRT Cell 8M (Catalog number 101-389-001, PacBio, Menlo Park, CA, USA) on the HiFi sequencing run.

### 2.8 Bioinformatic processing and analysis of 16S full-length sequences

RStudio environment along with DADA2 R package was used for quality control (QC) and taxonomical assignment of the PacBio generated V1-V9 16S raw reads (fastq files). First, low quality reads were filtered using QC DADA2 pipeline to perform quality filtering, end trimming, dereplication, error model specific to the dataset, ASV inference, and removal of chimeras [20]. Afterwards, taxonomical assignment was conducted using DADA2 Naive Bayesian Classifier algorithm using the SILVA version 138 16S full-length database [21]. Then, R packages phyloseq and ggplot 2 were used to generate relative abundance and 16S read count plots [22]. Comparisons between two cohorts at a single time point are calculated by Student’s t test. Comparisons between groups of more than two cohorts were calculated by ANOVA with Tukey’s (one-way) post-test. Comparisons between group variations were calculated by the PERMANOVA (permutational multivariate ANOVA) test.

### 2.9 Data sharing

The data have been deposited with links to BioProject accession number PRJNA1066774 in the NCBI BioProject database (https://www.ncbi.nlm.nih.gov/bioproject/).

## 3. RESULTS AND DISCUSSION

In this study we aimed to design an approach for sampling artwork for biological and genomic studies. We focused on centuries-old metalpoint and chalk drawings on paper that were acquired and utilized specifically for study. We first recorded high-resolution images before any sample collection. The high-resolution photography was then used to further define and describe the areas for sample collection. Since the upper corners of drawings tend to be handled most frequently by the artist, those areas were emphasized for sampling (areas A, B, and C). Further, since the center portion of the artwork is normally not handled post-production, this area was also chosen for sampling (areas D, E, and F). For the collection of samples, we compared double swabbing, core punching (1.2, 2, 4 and 6 mm), and M-Vac wet vacuuming. After samples were collected, gDNA was isolated using the QIAGEN DNeasy PowerSoil Pro Kit [23]. Finally, full length 16S (V1-V9) sequencing was carried out using the PacBio Sequel II/IIe system with SMRT Cell 8M on HiFi sequencing run. In addition to collecting samples from the artwork, we also isolated and sequenced DNA from negative controls, the scientists handling the artworks, and the art curators to define contaminants from the environment and others (Figures S1 and S2).

To determine if there were major differences in the DNA yields from the three sampling approaches, we first analyzed the total isolated DNA content from swabs, core punches, and M-Vac filters. Total DNA (ng/μL) obtained from swabs, pooled core punches (1.2, 2, 4 and 6 mm), or the M-Vac filters were compared. There were no significant differences between the yields obtained from the swabs and the pooled core punches; however, a significant increase in DNA yield was observed from the M-Vac filters (Figure 2A). Of note, when assessing the total read count estimates based on the full length (V1-V9) 16S rRNA gene copy number after quality filtering of the sequencing data, similar trends in DNA concentration were observed (Figure 2B).

**FIGURE 2.**
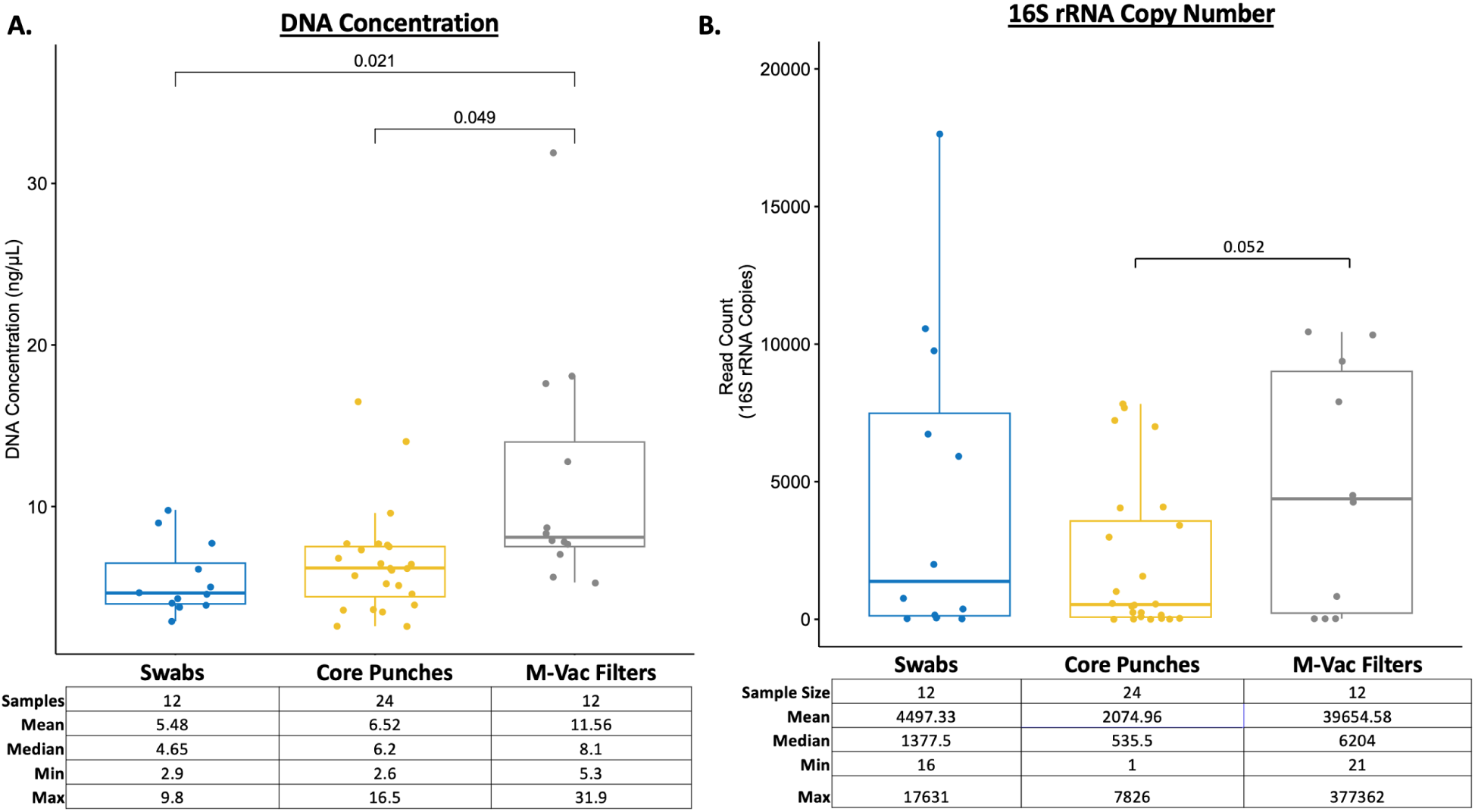
Total DNA content from swabs, core punches, and M-Vac filters. A) DNA concentration (ng/μL) of samples is based on the Nanodrop ND-1000 Spectrophotometer. T-test was performed by comparing the mean DNA concentration of the samples. We observed a significant (p-value <0.05) increase in DNA isolated from the M-Vac filters compared to swab samples or core punches. B) Total read count is estimated based on the full length (V1-V9) 16S rRNA gene copy number after quality filtering of the sequencing data. Similar trends to DNA concentration, an increase in the read count was observed from M-Vac filters.

We then aimed to determine the differences between the DNA yields obtained from the four different core punch sizes (1.2-, 2-, 4- and 6-mm). A t-test was performed using the 1.2-mm core punch as reference to compare against the mean read count of the other core punches at 2-, 4- and 6-mm sizes. The results showed that only the higher surface area of the 6-mm core punch led to significantly more bacterial DNA (Figure 3). As with the M-Vac wet vacuuming technique, the higher AS7 to AS10 values for the 4- and 6-mm core punches may produce irreparable damage to the artwork, as the recovered medium is destroyed during DNA isolation. Therefore, the core punch technique should only be used if preservation of the artwork is not necessary or when sampling can be conducted beneath the framed regions of the artworks.

**FIGURE 3.**
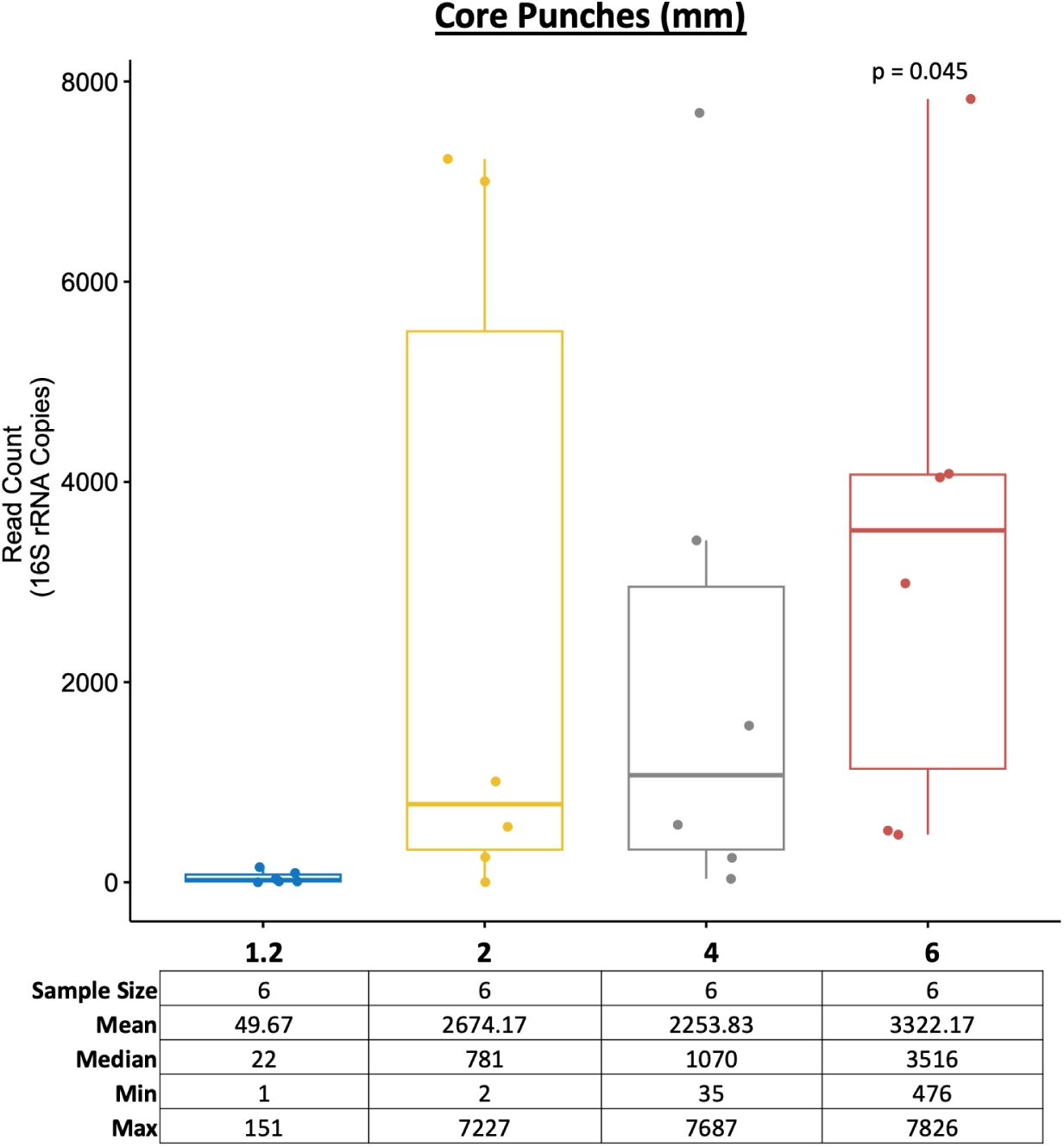
Core punches (6 mm) provides significantly (p-value <0.05) increased bacterial read count compared with smaller sizes. The t-test was performed using the 1.2-mm punch hole samples as reference to compare the mean read count of the other core punches (*i.e.*, 2-, 4- and 6- mm sizes). As expected, the amount of bacterial DNA obtained is positively correlated with the size of the core punches.

To determine if the location of the collected samples on the drawing affected the overall microbial readout obtained, we compared the total read counts for the full-length 16S rRNA obtained from the corners and the center of the artwork (Figure 1A). Comparisons between the corner and center samples for each collection technique (swab, core punch, and M-Vac filter) showed no significant difference of the mean read count (Figure 1B). These results suggest that while different collection approaches may improve the recovery of DNA, the microbial DNA content readout is similar irrespective of the localization within the artwork. This result was surprising since we expected more bacterial contamination in the corners and on the edges of the drawings. Furthermore, as noted later, there were no significant differences in the types of bacterial taxa found in the different locations, although the detection of diverse taxa was dependent on the sampling technique.

To define the bacterial profiles that are associated with each of the collection techniques, we determined the relative abundance of all bacteria identified in the full-length 16S sequencing data. A total of 150 operational taxonomic units (OTUs) were detected in the sequenced samples and are presented in an abundance plot (Figure 4). Interestingly, the number of OTUs varied based on the sample collection technique. Only 12% of the total OTUs observed were detected by all three sampling techniques. The wet vacuum technique detected 53% of the total OTUs observed, which was the highest among the three sampling techniques that were tested. Hence, these findings strongly indicate that sampling a larger surface area and retrieving sample from within the layers of the artworks will undoubtedly lead to the identification of more bacterial species. As shown by Alpha diversity index (Figure 6A), wet vacuum significantly yielded higher species richness compared to other sampling techniques.

**FIGURE 4.**
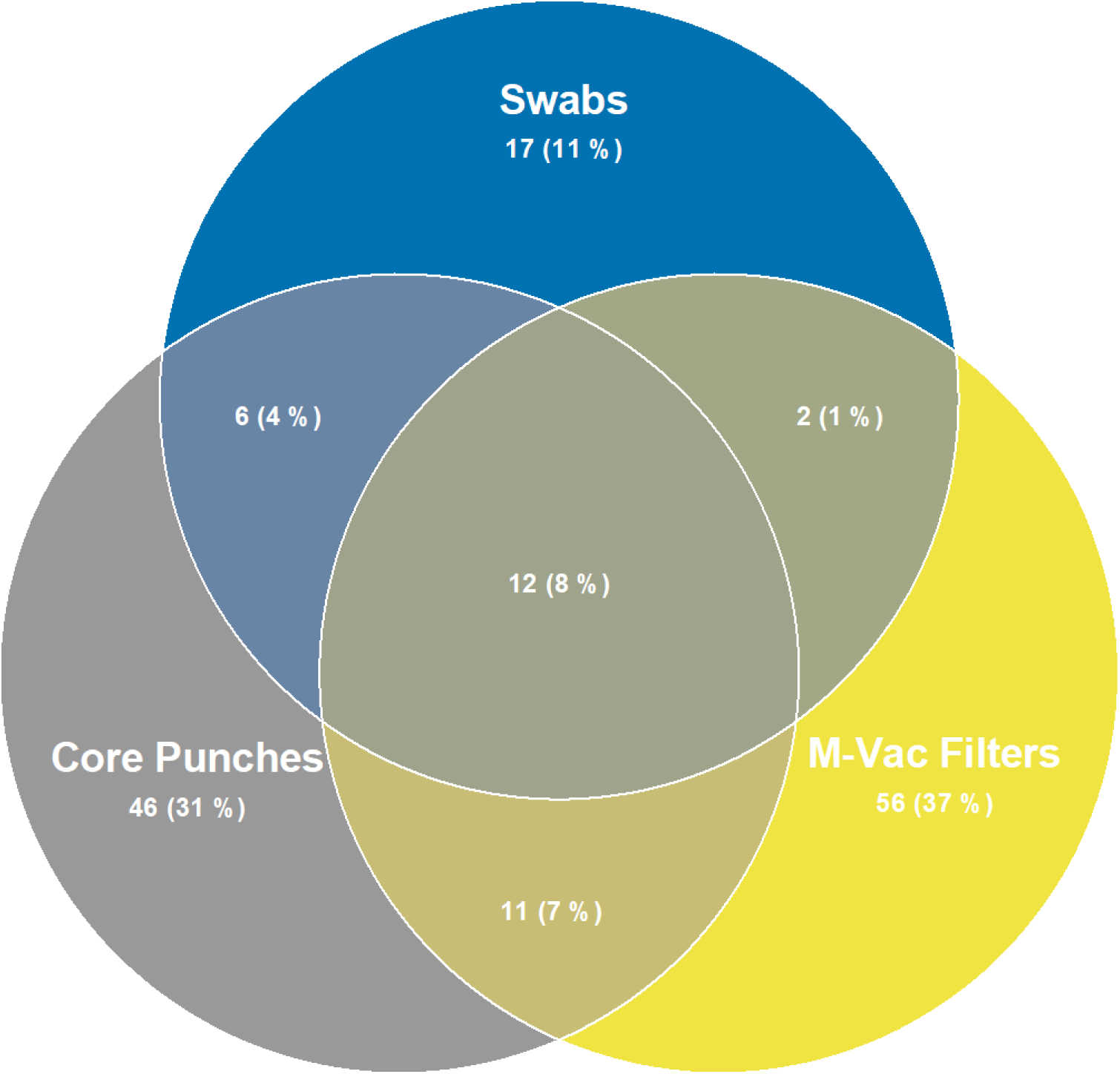
Comparing shared and unique operational taxonomic units (OTUs) for different sample collection techniques. A total of 150 OTUs were detected. Overall, 12 OTUs were shared among the collected sample types. Each sample collection techniques detected a different amount of OTUs. M-Vac filters (n= 81) and core punches (n= 75) detected more OTUs compared to swabs (n = 37).

Relative abundance of all bacteria (150 OTUs) identified in 16S full-length data showed differences in bacteria profiles based on sample type collected (Figure 5). The top 15 most abundant taxa identified in the samples collected from the different techniques ranged from environmental-associated to host-associated bacteria (Figures S3-S5). Of note, there were some observable differences in taxa associated with specific collection techniques, suggesting that each technique may provide access to different regions of the artwork and different biological readouts. Different species of bacteria were visibly abundant in the different sample collection techniques, for instance, *Turicibacter* sp. from swabs, *Bacillus* sp. from M-Vac filters, and *Anaerococcus* sp. from core punches. Of note, human commensal skin-associate bacteria, *Cutibacterium acnes* and *Staphylococcus hominis* [24], potentially originating from artwork handlers were abundant in M-Vac filters and core punches. In addition, these results may also suggest that combinatorial sample collection techniques may provide a more complete microbial profile of the tested artwork.

**FIGURE 5.**
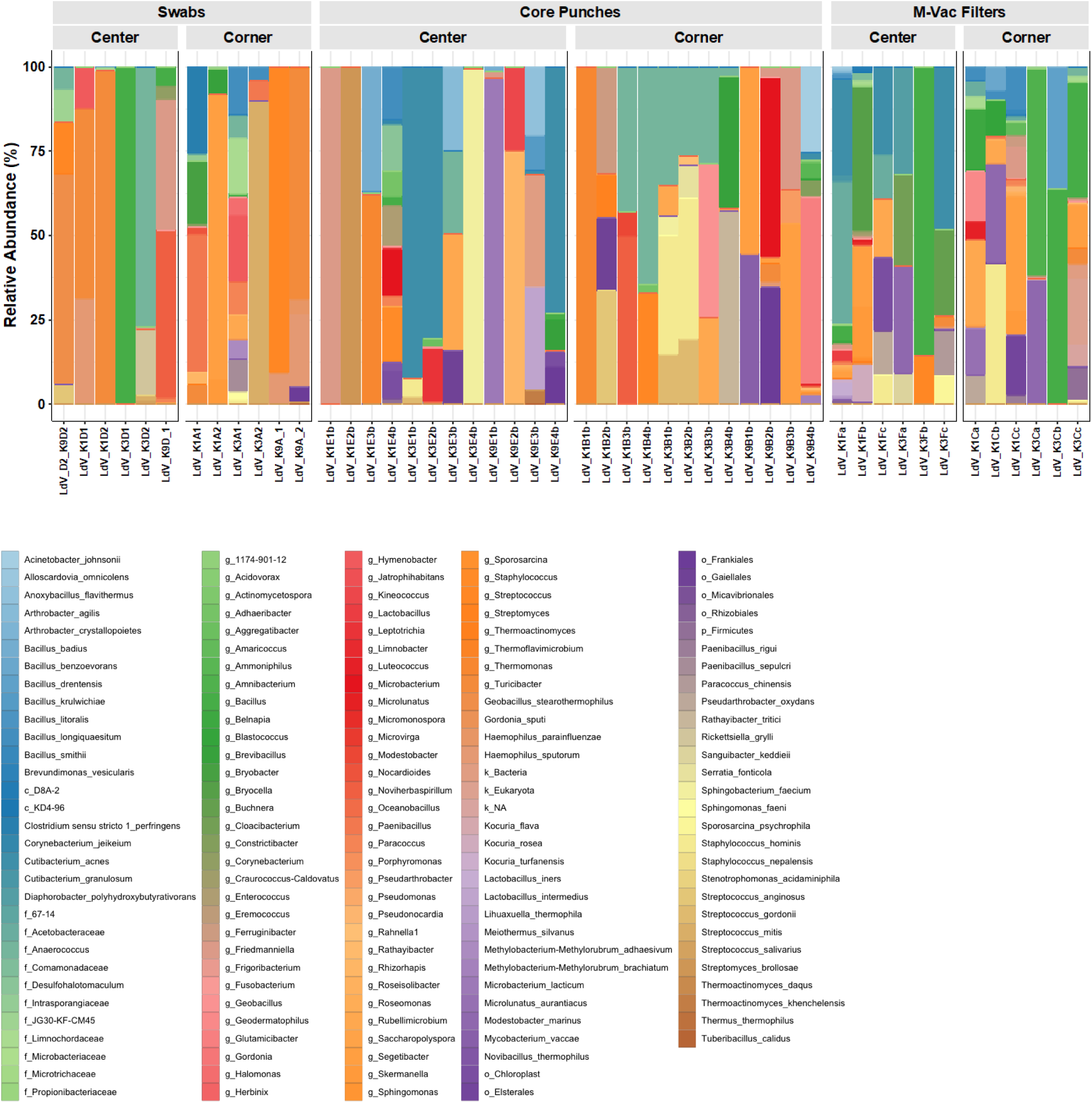
Different bacteria profiles identified based on sample collection techniques (swabs, core punches, and M-Vac filters). Relative abundance of all bacteria identified in 16S full-length data. Different sample collection techniques show the differences in bacterial community composition.

We then assessed the overall microbial diversity changes among the collection methods. The Alpha diversity index (Chao1 and Shannon) showed a significant (t test-based p-value <0.05) increase in species richness of samples collected using M-Vac filters compared with swabs and core punches, with no significant changes in diversity observed between the swabs and core punches (Figure 6A). Beta diversity, a measure of the variability in the community composition, showed that the overall microbiome of each of the collection techniques was similar based on sample clustering (Figure 6B). Using a PERMANOVA test, we aimed to define the association of microbial composition with variables such as sample collection techniques and the location of the sample collected. The test showed significant difference (R^2^ = 6% and p-value 0.01) in the artwork microbiome based on the sampling technique (M-Vac filters, core punches, and swabs), as observed in Figure 5 and Figure 6. However, the location of the sample collected within the artwork did not significantly contribute to differences in the artwork microbiome (R^2^ = 3% and p-value 0.06) (Figure 6C).

**FIGURE 6.**
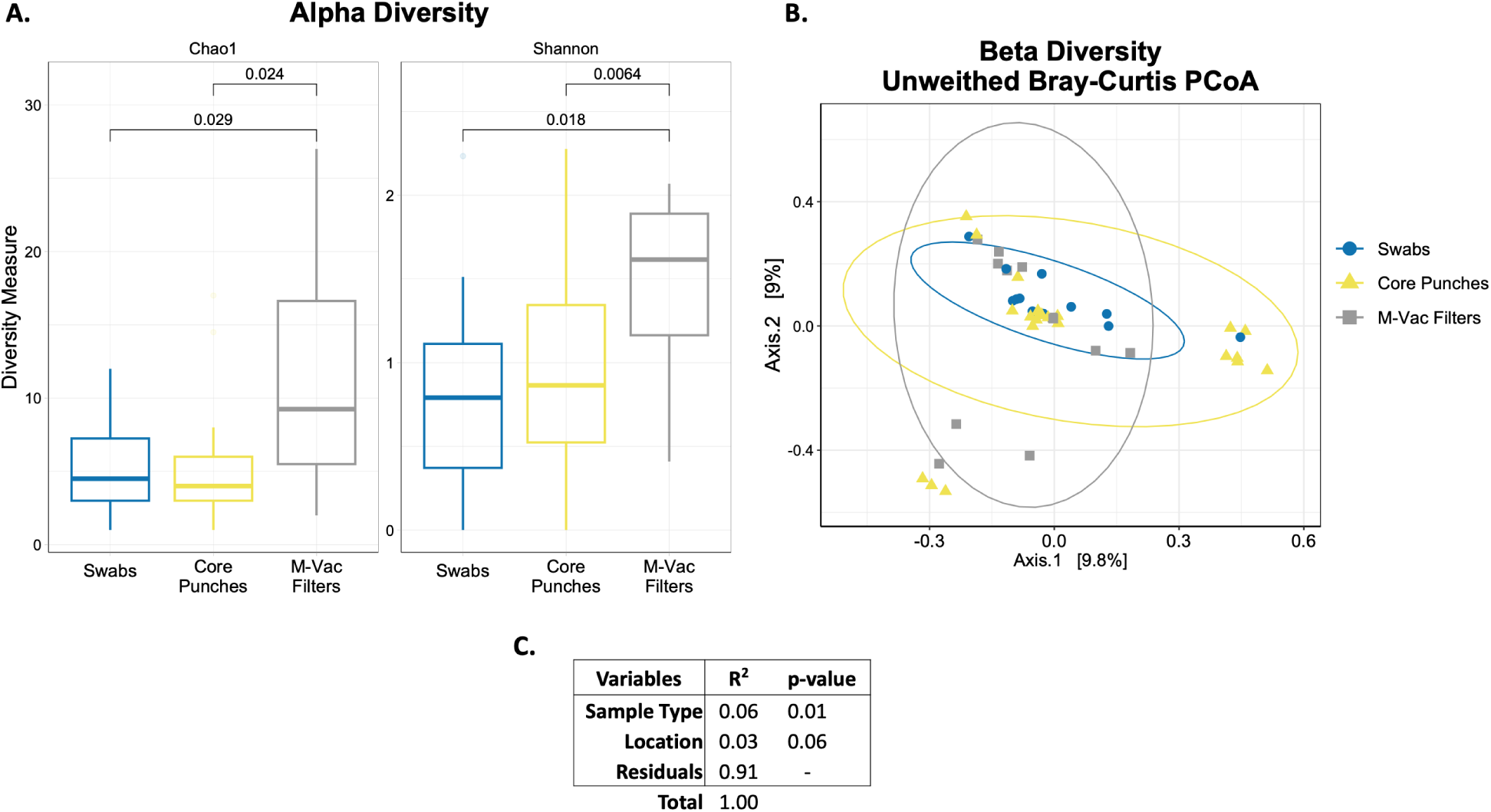
Diversity measures shows differential microbial composition between collection techniques. A) Alpha diversity index (Chao1 and Shannon) showed significant (t test p-value <0.05) increase in species richness of samples collected using the M-Vac filters compared to the other collection techniques. B) Beta diversity shares similarities in microbiome composition based on sample clustering. C) The PERMANOVA test shows significant difference (R^2^ = 6% and p-value 0.01) in the artwork microbiome based on the collection technique(swabs, core punches, and M-Vac filters). The location (triangles) of the sample artwork does not significantly contribute to the difference (R^2^ = 3% and p-value 0.06) to the artwork microbiome from the artworks.

## 4. CONCLUSIONS

In conclusion, this study shows the efficacy of various invasive and less invasive collection techniques to generate biological data from artwork. Here, we observed that the M-Vac wet vacuuming technique was able to recover consistently more biological material than the double- swabbing technique or the core punches. The wet vacuum approach was observed to alter the structural support of the artwork, the double-swab technique was observed to smear and remove some of the chalk from the red chalk drawings, and the core punches made obvious visible holes in the artworks. All three techniques were efficient in providing organic material for DNA isolation and sequencing. Of note, our data did not find any significant differences in the mean read count (microbial biomass) between the corners and the center of the artwork, suggesting that outside regions away from the main focus of the artworks may be feasible areas to sample to reduce damage to the artwork and obtain genetic material. Overall, our data show that double swabbing or small core punches provide an efficient route for sample collection for future studies of microbial material from artwork. However, it is still to be defined if these approaches can be used for isolation of human DNA in artwork. Since in most cases it is essential to utilize the most non-invasive, non-destructive sampling technique possible to preserve the integrity of the artwork being sampled, the double-swabbing technique described here should first be considered for sampling artwork for genomic analysis.

## ACKNOWLEDGMENTS

This work was carried out in the context of the Leonardo Da Vinci DNA Project. We thank Jesse Ausubel and Marguerite Mangin for their support and organizational efforts. We thank Angela Zimm and the late Fred Kline for their encouragement and assistance. Financial support was provided by the Richard Lounsbery Foundation. Full length 16S library preparation and sequencing was performed by the Institute for Genome Science, University of Maryland School of Medicine, Baltimore, MD, USA.

## CONFLICT OF INEREST STATEMENT

The authors have no conflicts of interest to declare.

Supplementary data encompasses six additional figures. Supplementary data to this article can be found online.

