## Supplementary material for "Sampling Techniques and Genomic Analysis of Biological Material from Artworks": Sampling Techniques_Roby_Supplemental Data.docx

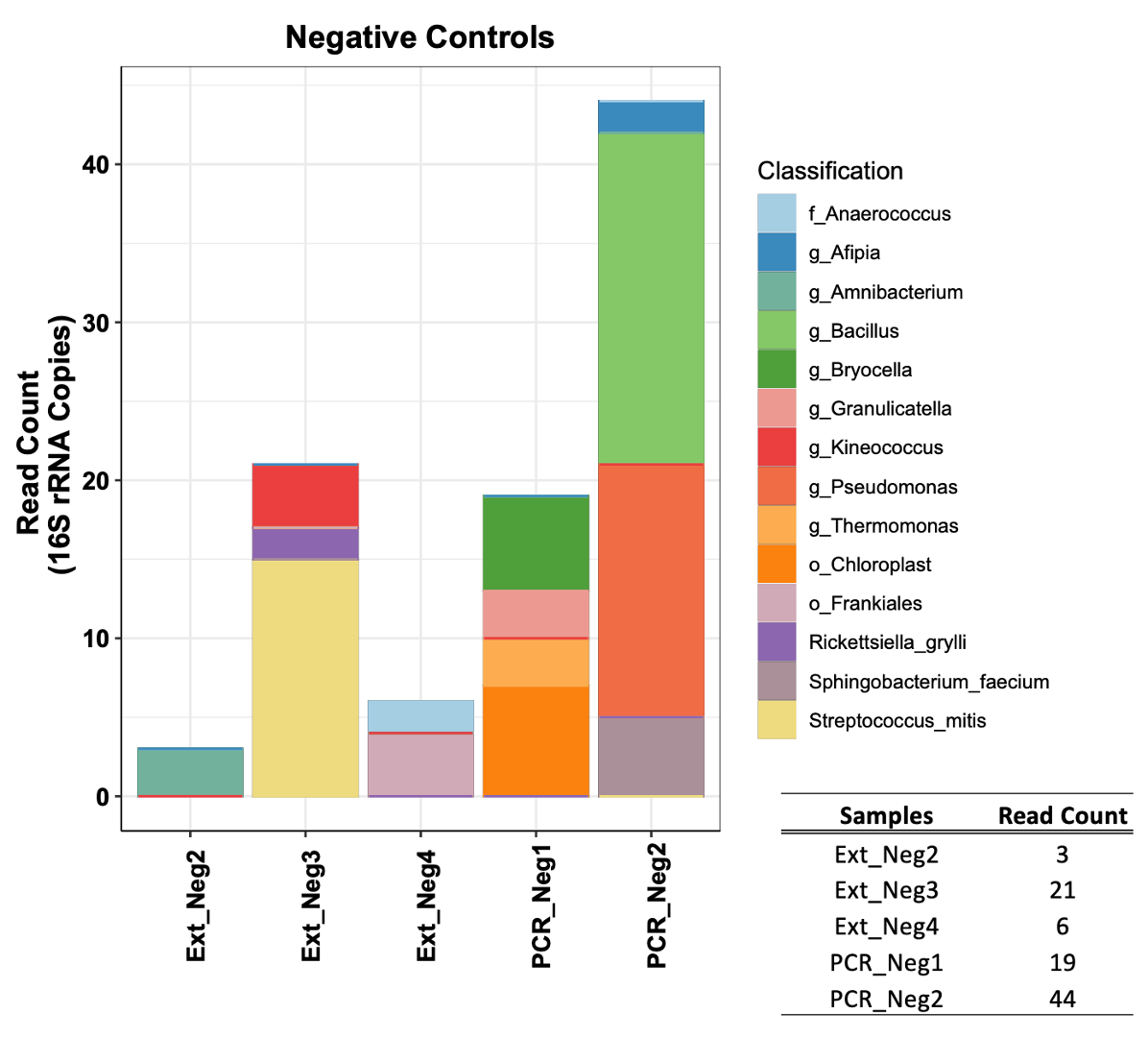

**Figure S1: Low contamination was identified in the extraction and PCR negative control samples.** Bacteria identified in the negative controls are associated with trace amounts of water- or reagent-associated bacteria taxa (e.g., *Pseudomonas* sp.) and human commensal bacteria (e.g., *Streptococcus mitis*).

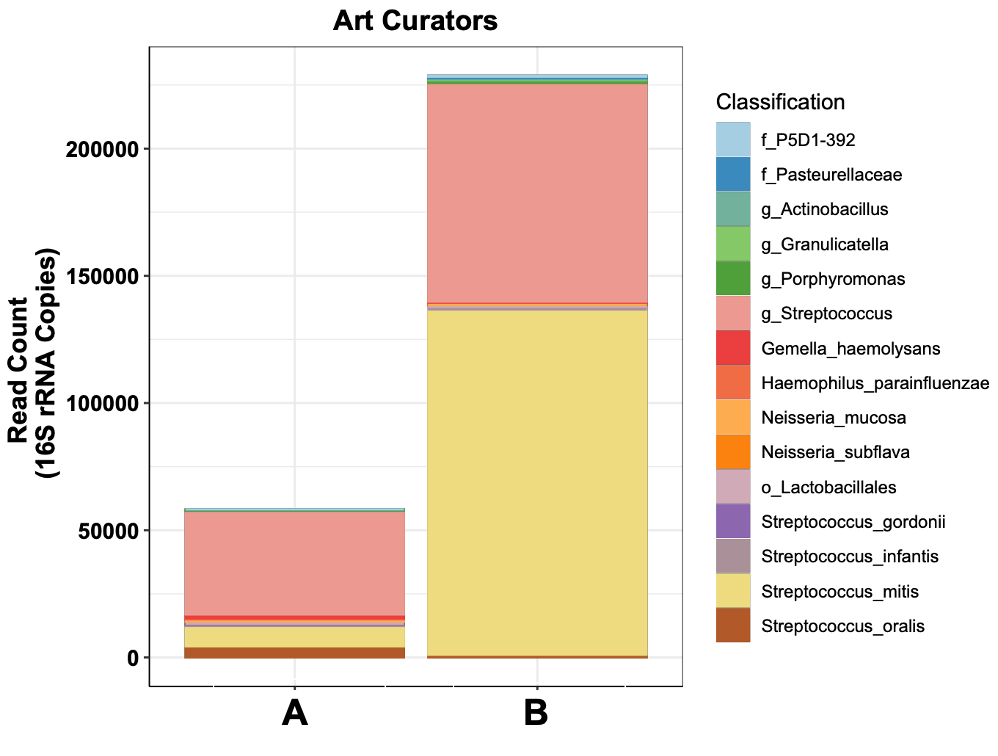

**Figure S2: Buccal swab control samples from the art curators to monitor potential human contamination,** The plot represents the total read count of the top 15 bacterial taxa identified in the samples. A total of 38 OTUs were detected in the art curator’s samples. Based on the full length 16S gene (V1-V9), a total of 58,821 reads were identified in sample **A** and 229,189 reads were identified in sample **B**. For both, host commensal bacteria *Streptococcus* sp. and *Streptococcus mitis* were most abundant. Human commensals *Streptococcus* and *Staphylococcus* sp. were detected in the centuries-old artwork samples (Figures S3-S4).

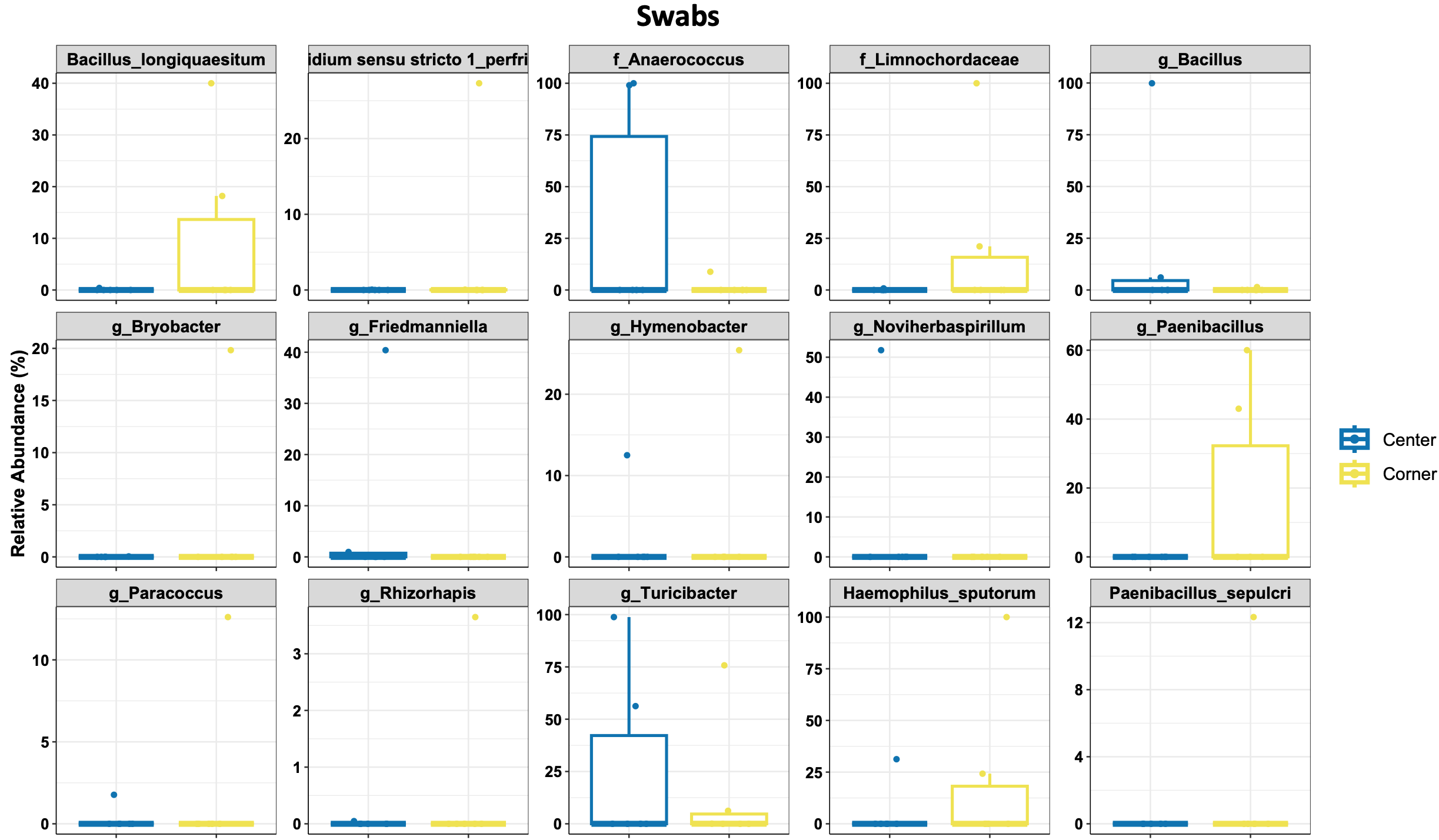

**Figure S3: Top 15 taxa identified in the swabs.** Significant differential abundant changes were identified in the corners and center. The center of the artwork had an abundance of *Anaerococcus* and *Turibacter* sp. The corners of the artwork had an abundance of environmental-associated taxa, *Bacillus longiquaesitum, Limnochordaceae,* and *Paenicacillus* sp. Host-associated bacteria, *Haemophilus sputorum*, was also abundant in the corners of the artwork.

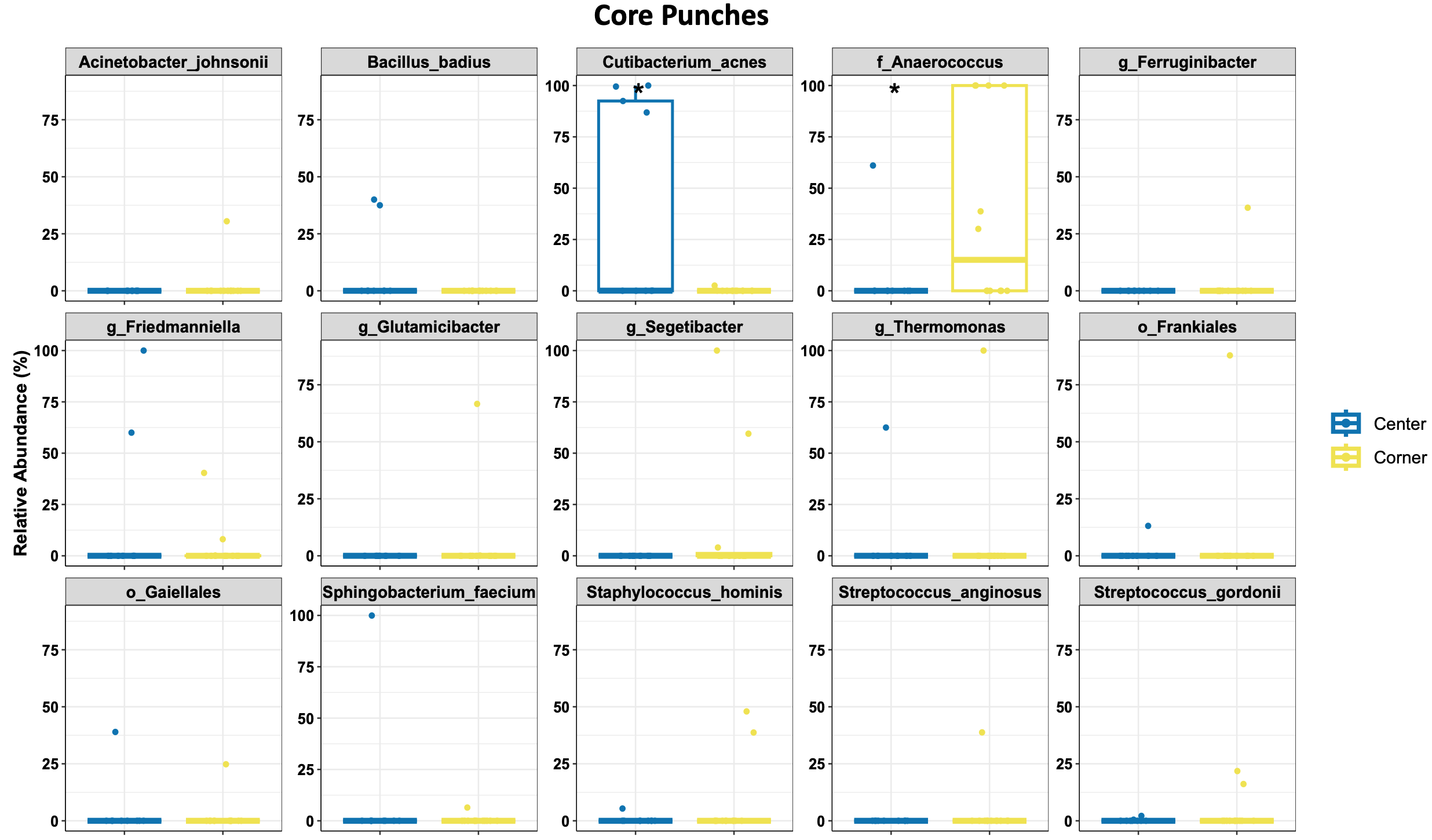

**Figure S4: Top 15 taxa identified in the core punches.** Significant differential abundant changes were identified in the corners and center. Commensal skin bacterium *Cutibacterium acnes* was significantly (p-value <0.05) detected in the center of the artwork. Host-commensal bacteria (commonly isolated from skin, human vagina, nasal cavity, oral cavity and feces), *Anaerococcus* sp., was significantly (p-value <0.05) detected in the corner of the artwork.

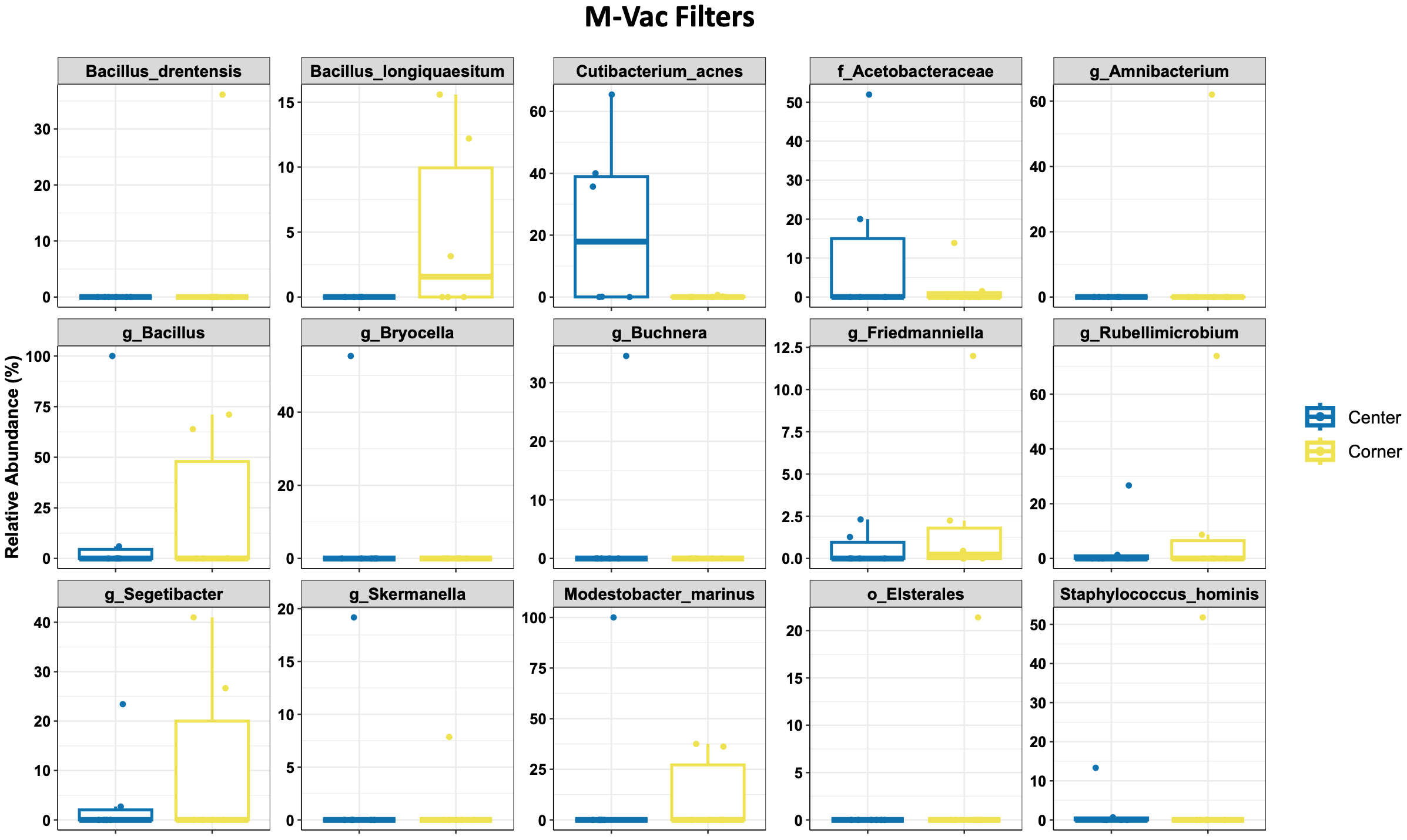

**Figure S5: Top 15 taxa identified in the M-Vac filters.** Differential abundant taxa were identified in the corners and center. For instance, *Cutibacterium acnes* (commensal skin bacteria) and Acetobacteraceae (family of strictly anaerobic bacteria that oxidizes ethanol to acetic acid) were abundant in the center. Abundant environmental-associated taxa identified in the corners of the artwork were *Bacillus longiquaesitum, Segetibacter* and *Modestobacter marinus*.
